# Genetic basis of the evolution of vertical bars in clownfishes

**DOI:** 10.1101/2025.03.14.643259

**Authors:** L. M. Fitzgerald, T. Latrille, A. Marcionetti, T. Gaboriau, D. A. Hartasánchez, N. Salamin

**Affiliations:** Department of Computational Biology, University of Lausanne, Lausanne, Switzerland

**Keywords:** Pomacentridae, Selection, Patterning, Coloration, Ancestral Reconstruction

## Abstract

Clownfish exhibit striking color patterns, characterized primarily by the presence of zero to three vertical white bars, along with three main colors: orange, white, and black. The common ancestor of clownfish likely possessed three vertical bars, with several instances of gains and losses occurring throughout clownfish evolutionary history over the past 10 million years. However, the evolutionary genomic mechanisms underlying the gain or loss of vertical bars remain unknown. In this study, we tested whether vertical bar transitions across the clownfish phylogeny were associated with changes in non-synonymous to synonymous substitution rates (*d*_N_*/d*_S_ values). Our analyses identified pigmentation-related genes that underwent changes in selective pressure, including *gch2, oca2*, and *vps11*, which are linked to melanophores, iridophores, and visual function. Additionally, *pmel*, a key melanogenesis gene, was found under positive selection, suggesting its role in shaping bar patterning. These results provide new insights into the genomic basis of coloration in clownfish, highlighting how selection and genetic variation influence phenotypic evolution.

## Introduction

Coloration in organisms can serve as a pivotal element in facilitating various ecological and evolutionary functions, such as signaling through inter- or intraspecific communication. Reef fish exhibit a wide range of color patterns, such as eyespots, vertical bars, and horizontal stripes, which can serve multiple purposes, including communication and predator-prey interactions (Salis et al., 2019a). For instance, colorful stripes in cleaner gobies (*Elacatinus* species) have been shown to facilitate communication in a cleaning mutualistic relationship, where the blue stripes of cleaner gobies serve to advertise their role to client fish, promoting cleaner-client interactions and deterring predators. In particular, yellow-striped gobies, while less conspicuous to predators, engage in cleaner-client interactions less frequently than their blue-striped counterparts, highlighting the evolution of coloration from predator deterrence to a more specialized signaling function (Lettieri and Streelman, 2010). Similarly, in butterflyfishes (*Chaetodon* species), stripes and eyespots have traditionally been linked to predator defense, such as misdirecting attacks or deterring predators. However, a comparative analysis suggests that these markings are evolutionarily dynamic and may also be shaped by ecological factors such as habitat type, sociality, and dietary complexity, rather than serving exclusively as anti-predator adaptations (Kelley et al., 2013). Thus, reef fish coloration plays a critical role not only in survival through predator avoidance but also in social behaviors, including territorial interactions and species recognition. The evolution of these patterns is likely driven by a complex interplay of selective pressures, highlighting the diverse functions of coloration in marine environments.

Within the reef fish family of *Pomacentridae*, the clownfish clade *Amphiprioninae* is a unique group of 28 species that form an obligate relationship with up to 10 species of sea anemones (genera *Entacmaea, Stichodactyla*, and *Radianthus*), and this association was suggested to have likely triggered their adaptive radiation (Litsios et al., 2012). Clownfish have a relatively simple pattern of three main colors (black, white, and orange) and zero to three vertical white bars. Some species, mostly from the skunk species complex, have a horizontal stripe across the top of their head and body. The monophyletic nature of the clownfish clade allows for evolutionary replicates, where similar color patterns have evolved multiple times across species. This provides an opportunity to investigate the ecological and molecular mechanisms underlying parallel evolution in color patterns. Recently, it was found that the coloration of clownfish is significantly linked to their host sea anemone (Gaboriau et al., 2024). Individuals hosted by *Radianthus* sea anemones tend to be more orange/yellow, those hosted by *Entacmaea* are darker orange/red, and those that are generalists (inhabiting in more than three anemone species) are darker with contrasting vertical bars.

In the literature, several hypotheses have been proposed about the role of color patterns in clownfish (Hayashi et al., 2022). One is species recognition, which serves as a form of communication and dominance between individuals with different numbers of white bars. In juvenile *Amphiprion ocellaris*, individuals were more likely to attack other individuals with different numbers of vertical bars than those with the same number of bars (Hayashi et al., 2024). Additionally, observations of interspecies cohabitation are rare, but when it occurs, a consistent absence of identical bar configurations among these coexisting species exists. For example, individuals from *A. clarkii* (two or three bars) can be found with *A. perideraion* individuals (one bar). Similarly, individuals of *A. biaculeatus* (three bars) can coexist with *A. melanopus* individuals (one bar) (Camp et al., 2016). Another suggested role of color patterns is protection from predators, as the contrasting colors and white bars help hide the fish silhouette within the sea anemone tentacles (Salis et al., 2018). Finally, a function of aposematic signaling, in which clownfish use their striking colors to advertise the toxicity of their host sea anemone has also been proposed (Merilaita and Kelley, 2018). However, these hypotheses are not conclusive to the function of coloration and bar patterning in clownfish.

Understanding the function of clownfish color patterns provides important context for investigating how these traits evolved. Ancestral state reconstruction can help clarify whether certain patterns were historically linked to ecological pressures such as species recognition, predator avoidance, or aposematic signaling. One first study (Merilaita and Kelley, 2018), used a maximum likelihood estimation to infer the ancestral vertical bar patterns treating the number of vertical bars as a continuous trait. They showed that the ancestral clownfish had on average 2.73 vertical bars and evolutionary transitions most likely led to a bar loss rather than a gain. The loss of vertical bars was not exclusive to the clownfish clade but occurred several times across different parts of the phylogenetic tree of the *Pomacentridae* family. In contrast, a second study (Salis et al., 2018) used stochastic character mapping, which is a Bayesian approach that incorporates uncertainty in the character evolution. The transition rates between number of vertical bars were estimated using a likelihood based model where they compared equal-rates to unequal-rates transition models, focusing on the loss of vertical bars. They showed that the common ancestor of clownfish likely had three vertical bars approximately 12 million years ago. The color patterns seen in extant species could thus result from losing their vertical bars caudal-rostrally. For instance, some species have more vertical bars as juveniles and lose them caudal-rostrally as they age. In addition to their different approaches, both of these studies used a phylogeny based on seven nuclear markers (Litsios et al., 2014), which has been recently updated using whole genome data of the 28 species (Gaboriau et al., 2024).

To confirm these evolutionary patterns, it is essential to investigate the molecular mechanisms underlying color production and pattern formation. Colors can be produced by organisms through structural coloration and pigmentation (Orteu and Jiggins, 2020). Structural colors are produced by interference diffraction or scattering of light layers, while biological pigments selectively absorb and reflect different wavelengths of light (Booth, 1990). In teleost fish, there are three main types of pigment cells (also known as chromatophores), namely, melanophores, xanthophores, and iridophores, as well as less common types such as leucophores, erythrophores, and cyanophores (Salis et al., 2019b). Several studies have linked these chromatophores to specific genes in fish (Lorin et al., 2018; Salis et al., 2021; Moore et al., 2023). These studies suggest that pigmentation is influenced by a combination of genetic factors, with some genes having more significant effects than others. While the genetic basis of color pattern formation is complex, involving multiple loci, the exact nature of the genetic architecture –whether polygenic or governed by supergenes– remains unclear.

Understanding the regulatory mechanisms that govern these genetic factors is critical, as gene expression plays a fundamental role in shaping phenotypic plasticity, particularly in relation to clownfish color patterns. For example, in *A. percula*, thyroid hormones (THs) regulate the expression of *duox*, a gene involved in pigment cell development. Increased *duox* expression leads to faster white bar formation, a process influenced by their anemone host species. Juveniles form white bars faster when associated with *Stichodactyla* compared to *Radianthus* anemones (Salis et al., 2021). In juvenile *A. ocellaris*, THs also modulate opsin gene expression, affecting visual perception (Roux et al., 2023). These findings suggest that gene expression contributes to both white bar formation and visual adaptation.

While gene expression governs immediate phenotypic responses, protein-coding sequences are subject to evolutionary pressures that shape long-term trait evolution. Genes associated with pigmentation and visual perception may experience different selective regimes, depending on ecological or behavioral constraints. For example, genes involved in visual perception could be under positive selection in species where visual acuity plays a crucial role in predator detection or mate choice. Similarly, pigmentation genes may evolve under different selective pressures depending on the ecological or social context. To evaluate selective pressures acting on protein-coding DNA sequences, a typical proxy is the non-synonymous substitution rate (*d*_N_), which is compared to the synonymous substitutions (*d*_S_), calculated as a ratio called *d*_N_*/d*_S_ or *ω* (Goldman and Yang, 1994; Muse and Gaut, 1994). In this framework, methods such as RELAX (Wertheim et al., 2015) detects shifts in the strength of natural selection by comparing whether selection has been relaxed or intensified across a phylogeny, employing a binary framework that contrasts foreground versus background branches of discrete traits. In contrast to a binary framework, the change in selective pressures acting on genes can be modeled as changing along branches of the phylogenetic tree (Seo et al., 2004). This modeling provides a more nuanced understanding of selection, and the Bayesian integrative framework allows for the detection of changes in selective pressures associated with other traits (Lartillot and Poujol, 2011; Latrille et al., 2021). Regarding rates of evolution compatible with episodes of positive selection, CODEML (PAML) (Yang, 2007) seeks *d*_N_*/d*_S_ values greater than 1 for some specific sites of the protein-coding genes. In *Pomacentridae*, a study examining selection in six opsin genes within the family found all but one visual opsin gene under positive selection. Additionally, they found cases of divergent, parallel, convergent, and reversed evolution across the damselfish family, highlighting the importance of visual acuity in the group (Stieb et al., 2017). Given the link between TH-mediated gene expression and pigmentation, genes regulating white bar formation in clownfish may be also evolving under specific selective pressures. A study identified 128 pigmentation-related genes in clownfish, categorized by pigment cell type (Salis et al., 2021), providing candidate genes for further evolutionary analysis.

We hypothesize that specific genes under directional selection or having undergone changes in selective pressure are related to the gain or loss of white vertical bars in clownfishes. Genes essential for maintaining bars are likely under stronger purifying selection, whereas those associated with bar loss may exhibit relaxed selection. Conversely, if vertical bars provide an adaptive advantage– such as species recognition, genes involved in their formation may experience directional selection. By examining changes in *ω* values across pigmentation genes and mapping these changes onto the phylogeny, we aim to identify signatures of directional selection or changes in selection pressure associated with vertical bar evolution and clownfish coloration. By integrating phylogenetic and genomic approaches, this study provides insights into the molecular mechanisms underlying vertical bar evolution. These findings will contribute to a broader understanding of how selection shapes phenotypic diversity and adaptation in marine organisms.

## Materials & Methods

### Data

Throughout the study, we tested two sets of genes. The first dataset was curated from the 128 pigmentation genes, categorized by pigment cell, previously identified by Salis et al. (2021). We identified orthologous genes present in all clownfish species through reciprocal best BLAST (Altschul et al., 1990) to *A. frenatus* (Marcionetti et al., 2018), reducing the initial list to the 79 “Color Genes” used in this study (Supplementary Table 1). We also wanted to test if there were additional genes that could be related to color or pigmentation that were not found in the initial dataset so we created an “All Genes” dataset (which include the 79 Color Genes) and corresponds to 18,390 protein-coding genes shared across the updated clownfish phylogeny Gaboriau et al. (2024). We classified the number of vertical bars in adults as none, one, two, or three from Salis et al. (2018) classification (Supplementary Table 2). We did not include polymorphic traits for *A. clarkii* or *A. polymnus*, but kept the assigned number from Salis et al. (2018) that classified both species as having two vertical bars.

### Ancestral state reconstruction and branch categorization

We reconstructed the ancestral state of the number of vertical bars as adults using the R package*CorHMM* (v2.8) (Beaulieu et al., 2013). We tested different transition matrices corresponding to six evolutionary models: (1) all rates differ, (2) symmetric, (3) equal rates, (4) biological all rates differ, (5) biological symmetric, and (6) biological equal rates (Supplementary Table 3). The biological models (models 4-6) ensured logical progressions of the trait evolution (e.g., no transition from no bars to two bars without passing through one bar). The best model was selected based on the Akaike Information Criterion (AIC) (Akaike, 1974). We then used the R package *phytools* (v2.1-1) (Revell, 2012) to generate 100 stochastic maps under the best model. The summary of these 100 stochastic maps is shown in Figure 1. For each map, we categorized every branch as having gained, lost, or retained the number of bars between the parent and child nodes corresponding to the highest weight (Supplementary Figure 1, Filtered output). Weight differences for each branch were calculated by subtracting the loss weight from the gain weight.

**Figure 1:**
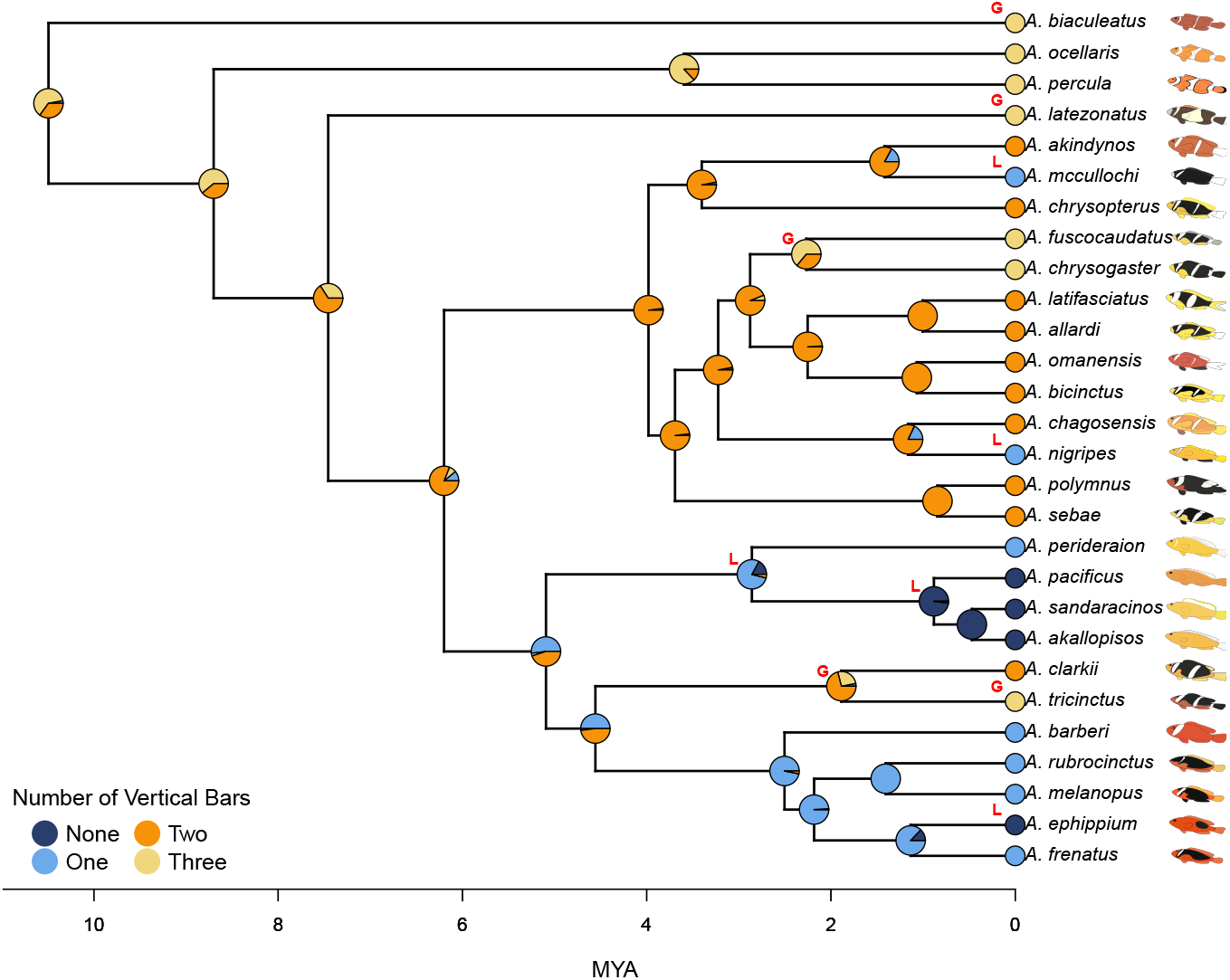
The summary stochastic map of the reconstructed number of vertical bars with the four states: None, One, Two, and Three. Pies at each node represent the marginal probability of each ancestral number of bars. “G” labels to the right of the node refer to branches categorized as Gain and “L” as Loss.

### Detection of differential selective pressure associated with the evolution of bar number

To test for an association between the gain or loss of bars and changes in selective pressures, we relied on the *d*_N_*/d*_S_ ratio, which can be denoted as *ω* at the gene level. The values of *ω* signify different forms of selection: *ω <* 1 indicates negative (purifying) selection, preserving protein function; *ω >* 1 suggests positive selection, potentially driving functional divergence; and *ω* = 1 indicates neutral evolution (Yang et al., 2000). Relaxed selection, where purifying constraints are reduced, can contribute to greater phenotypic variation (Lahti et al., 2009). We estimated *ω* changes along the clownfish phylogenetic tree using a Bayesian approach implemented in the *BayesCode* software (Latrille et al., 2021) using the following commands:

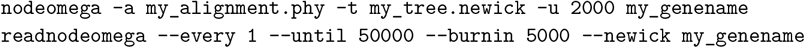

In *BayesCode*, the prior of *ω* along the tree is simulated as a Brownian process, and its value changes at each node of the tree, or, in other words, it is increasing or decreasing along each branch of the tree. The topology of the tree was fixed to correspond to the clownfish species tree of Gaboriau et al. (2024), while node ages and changes in mutation rate along the tree were jointly estimated by *BayesCode* (Latrille et al., 2021). We then obtained posterior estimates of Δ*ω*, representing for each branch the changes in *ω* along this branch. Altogether, we obtained one posterior Δ*ω* per branch and gene. In our analyses, a positive Δ*ω* implies a increased *ω* value on the focal branch, indicating some relaxation of purifying selection or increase of positive selection, which both indicate that more changes were occurring at the molecular level. In contrast, a negative Δ*ω* could suggest a tightening of selection that limits changes at the molecular level because the estimated *ω* is smaller in the most recent branch. For each gene, to test whether changes in selective pressure along branches of the tree were associated with the loss or gain of bars, we used a linear model to predict Δ*ω* as a function of bar changes (loss, same, gain). We obtained one value of Δ*ω* per branch, whereas the number of vertical bars changed depending on each stochastic map of our ancestral reconstruction (see previous section) (Supplementary Figure 1). Therefore, we used a weighted regression using the proportion of stochastic maps resulting in bar changes (loss, same, gain) as the independent variable. The significance of the weighted regression was assessed as a *p-*value for each gene. Across the genes tested, we corrected for multiple testing using the *p*.*adjust* function and the Benjamini-Hochberg method implemented in the R package *stats* (v4.2.3) (R Core Team, 2023). Significantly enriched Gene Ontology (GO) terms (http://geneontology.org/) of the significant genes were identified using Fisher’s exact test in the topGO package (Alexa and Rahnenführer, 2009).

### Testing for positive selection

To investigate the presence of genes under positive selection linked with the gain or loss of vertical bars, we used the branch-site model H1 (model = 2, fix omega = estimate) compared to a neutral model H0 (model = 2, fix omega =1) as implemented in CODEML (PAML, v.4.9) (Yang, 2007) across the “All Genes” dataset. We categorized branches (gain or loss) from the ancestral state reconstruction as foreground branches in the branch-site model. The model was run separately for the two types of transitions (i.e., gain or loss; Supplementary Figure 2). We followed the recommendations from PAML, in which CODEML was run three times for the null and the alternative hypothesis with the initial branch length ignored to allow for three randomized runs of the optimization process to increase the chances of reaching the optimum. The highest likelihood for each test run was saved for the likelihood ratio test (df=1). We extracted both the Bayes empirical Bayes (BEB) (Yang et al., 2005) and Naive empirical Bayes (NEB) (Nielsen and Yang, 1998) values from the CODEML output. Then we corrected the *p-*values across all genes for multiple testing using the *p*.*adjust* function and the Benjamini-Hochberg method implemented in the R package *stats* (v4.2.3) (R Core Team, 2023). Lastly, significantly enriched GO terms (http://geneontology.org/) of the significant genes were identified using Fisher’s exact test in the topGO package (Alexa and Rahnenführer, 2009).

**Figure 2:**
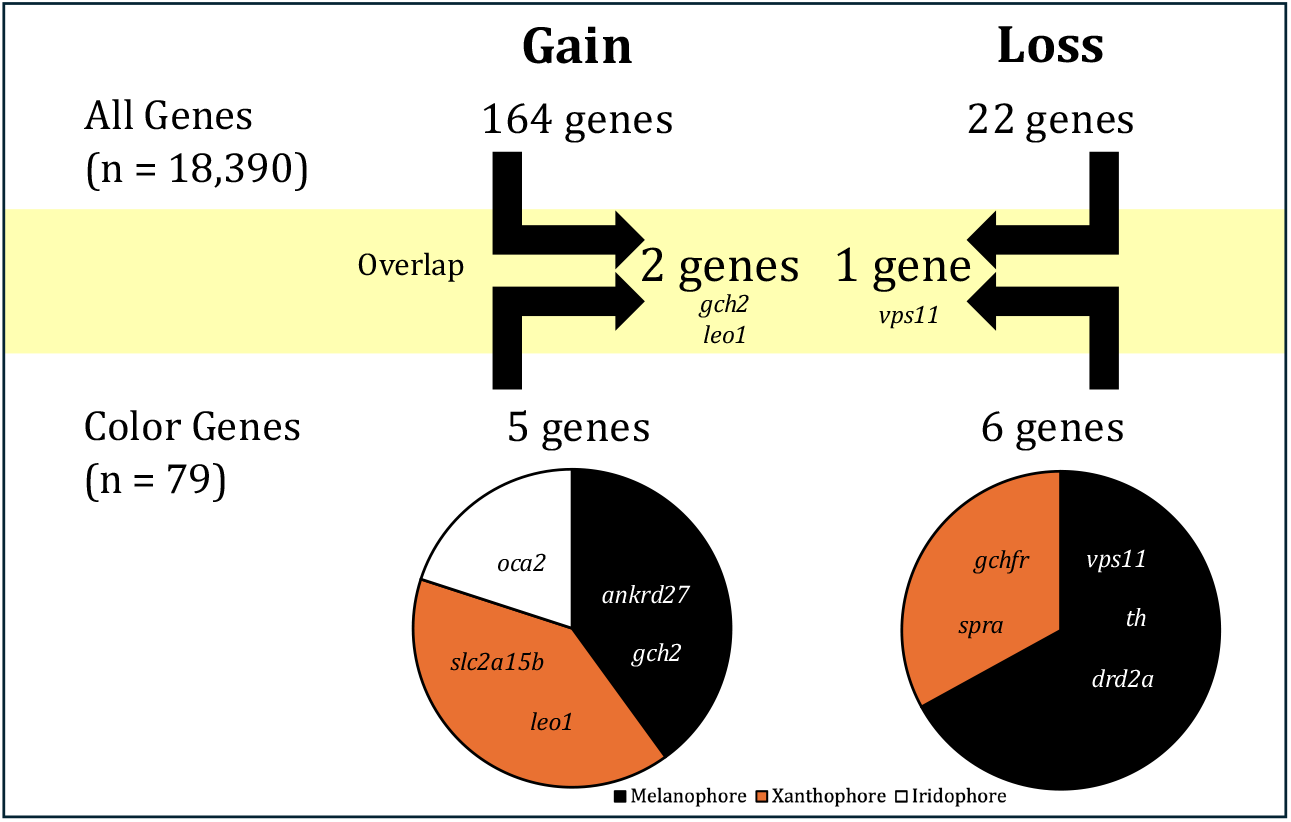
Summary of the number and name of significant color genes found in each dataset (All Genes and Color Genes) for gain and loss and their pigment cell type.

## Results

### Repeated gain and loss of vertical bars across the clownfish phylogeny

The number of vertical bars at each node of the phylogenetic tree was inferred using ancestral state reconstruction. The best model was selected based on the lowest Akaike Information Criterion (AIC) value, and corresponded to model 6 (AIC = 59.4), the biological equal rates model. This model assumed equal transition rates between any consecutive state (e.g., one to none, or two to three) and did not allow transitions between non-consecutive states (e.g., one to three). We summarized the 100 replicates of stochastic mapping by recording the number of vertical bars inferred at each node and estimating the probability of each state (Figure 1). The root node of the clownfish phylogenetic tree was inferred to have three vertical bars with probability = 0.62 (two bars: probability = 0.35; one bar: probability = 0.03). This number then evolved through gains and losses of bars within the main clownfish clades. We identified two internal branches (ancestors to *A. tricinctus* and *A. fuscocaudautus*) and three terminal branches (*A. biaculeatus, A. latezonatus, A. tricinctus*) as gain. In addition, two internal branches (ancestors to *A. perideraion* and *A. pacificus*) and three terminal branches (*A. mccullochi, A. nigripes*, and *A. ephippium*) were identified as loss. Most of the nodes are categorized as same (Supplementary Figure 1). Throughout the phylogeny, we have evidence for repeated cases of loss and gain of vertical bars. Two consecutive losses leading to the skunk complex and two consecutive gains leading to *A. tricinctus* might suggest an underlying adaptive process.

### Genes associated with gain or loss of vertical bars

Out of the 18,390 orthologous genes used in our study, we found 164 genes showing patterns of significant differential selection, Δ*ω* associated with vertical bar gain, while 22 genes were associated with vertical bar loss (Supplementary Table 4). We also obtained average *ω* values and average *ω* for gain or loss for each significant gene (Supplementary Table 5). There were no GO terms associated with coloration or vision in the significant genes associated to vertical bar gain (n=164) or loss (n=22) (Supplementary Table 6). Within the significant genes associated with bar gain, we found two coloration genes, and for bar loss, one coloration gene (Figure 2). Testing only within the “Color Genes” dataset, we found five genes under differential selection linked to vertical bar gain, and six genes under differential selection linked with vertical bar loss (Figure 2). Each of these genes is predominantly expressed in one pigment cell type: melanophores (black), xanthophores (orange), or iridophores (white). Genes associated with the gain of bars represent all three types of pigment cells, whereas only melanophores and iridophores are represented among genes linked to bar loss. Within the five genes related to vertical bar gain (*ankrd27, ω* = 0.40; *gch2, ω* = 0.16; *leo1, ω* = 0.09; *oca2, ω* = 0.05; and *slc2a15b, ω* = 0.06) (Supplementary Table 5) we found different branches driving the Δ*ω* in the significant coloration genes (Figure 3). Similarly, within the six coloration genes significantly related to vertical bar loss (*adam17b, ω* = 0.20; *drd2a, ω* = 0.11; *gchfr, ω* = 0.06; *spra, ω* = 0.22; *th, ω* = 0.28; and *vps11, ω* = 0.20) (Supplementary Table 5), we also found different branches driving the Δ*ω* in the significant coloration genes (Figure 4).

**Figure 3:**
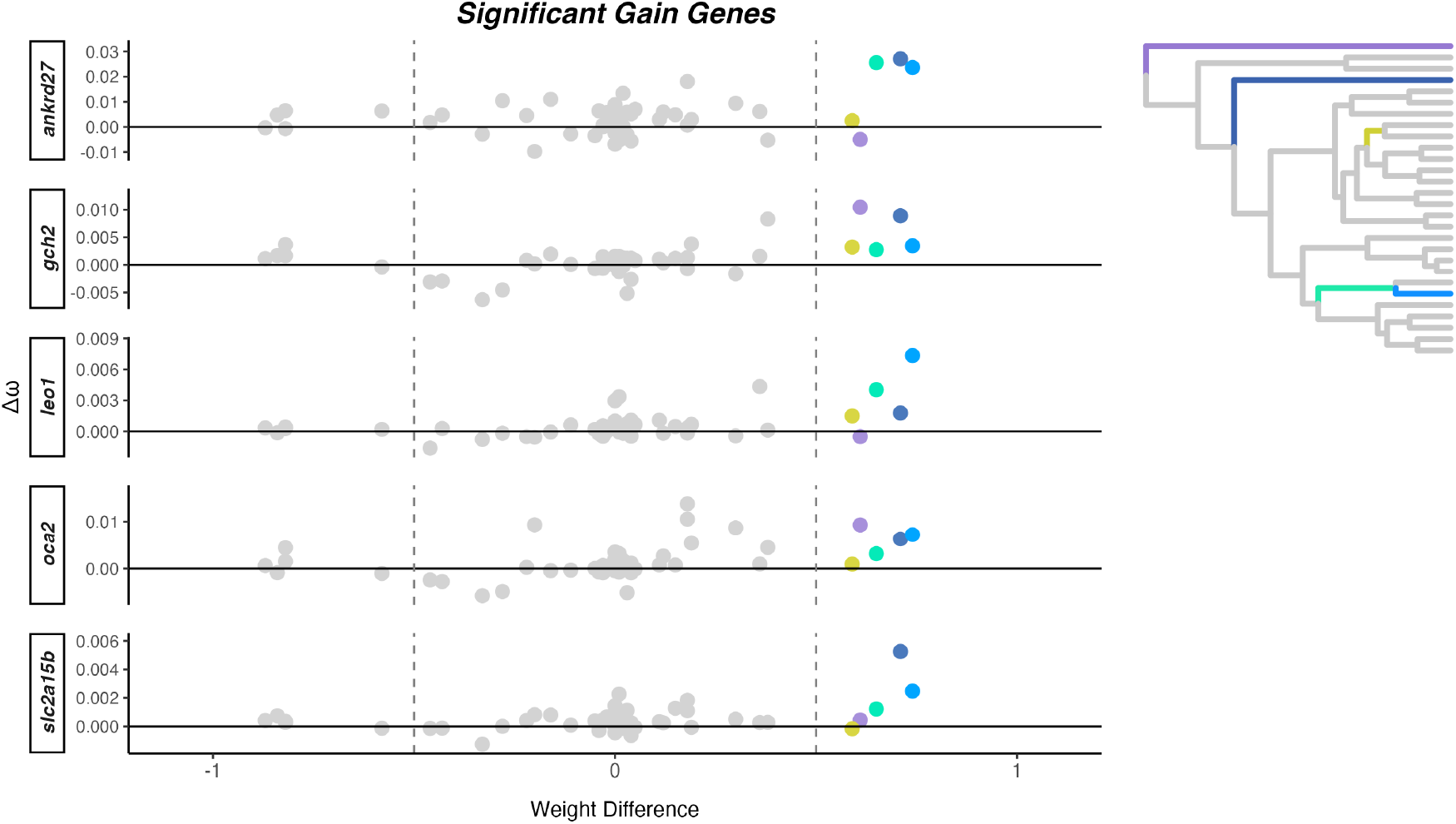
Δ*ω* and weight difference for five color genes significantly associated with vertical bar gain. Each dot represents a branch on the phylogenetic tree (n = 54), with colored dots representing branches that are categorized as gain, colored in the phylogenetic tree (upper right corner; a larger version is in Supplementary Figure 2).

**Figure 4:**
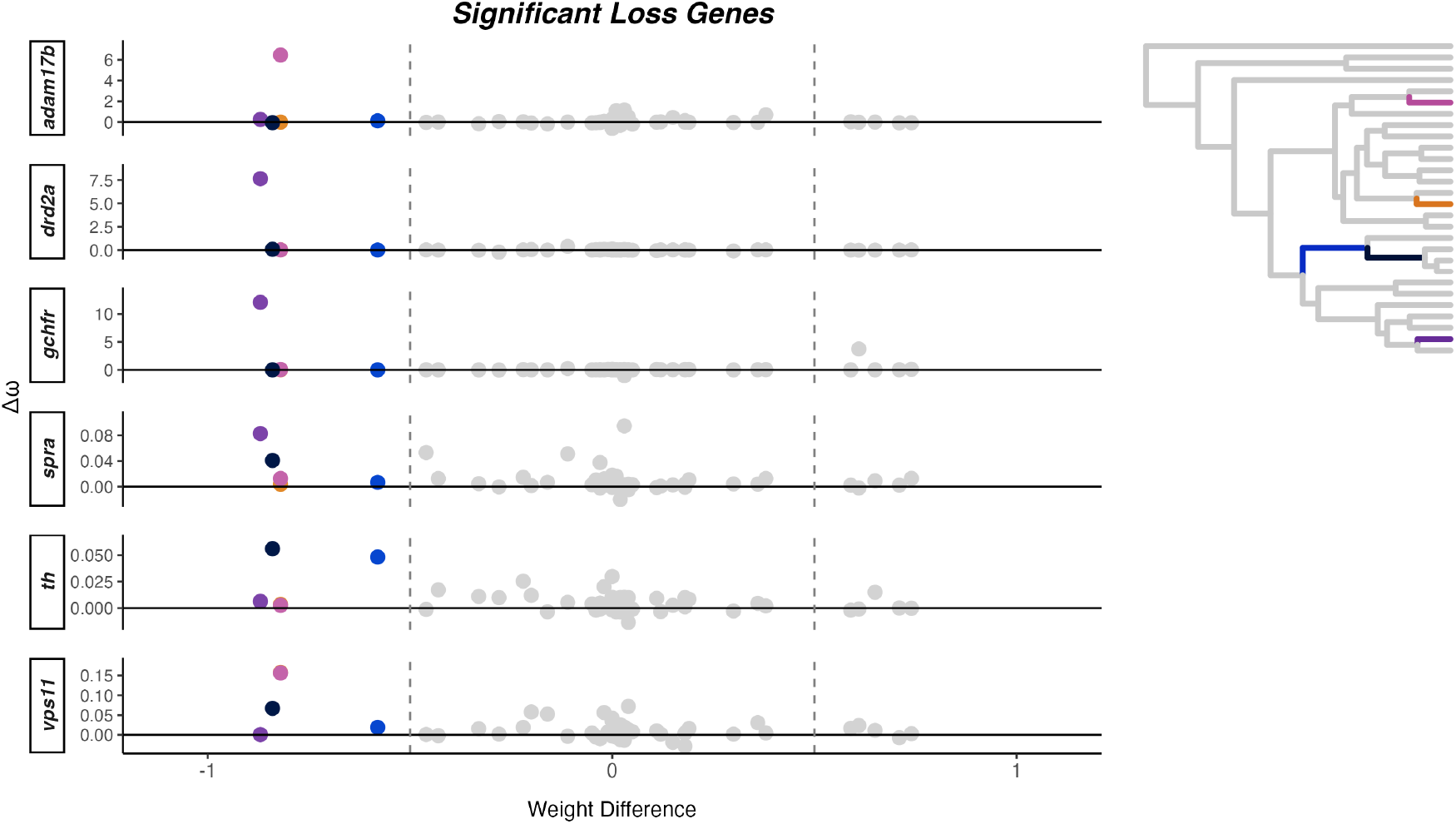
Δ*ω* and weight difference for six color genes significantly associated with vertical bar loss. Each dot represents a branch on the phylogenetic tree (n = 54), with colored dots representing branches that are categorized as loss, colored in the phylogenetic tree (upper right corner; a larger version is in Supplementary Figure 2).

### Positively selected genes related to vertical bar gain

Across the entire dataset (18,390 genes), we found 216 significant genes under positive selection associated with vertical bar gain, and 67 genes under positive selection with vertical bar loss (Supplementary Table 4). We also calculated the average foreground *ω* values for each significant gene (Supplementary Table 5). There were no GO terms associated with coloration or vision in the significant genes related to vertical bar gain (n=216) or loss (n=67) (Supplementary Table 6). Among the 216 genes related to vertical bar gain, two coloration genes were found, *saiyan (si:ch211-256m1*.*8)* and *pmela*. Although both Bayes empirical Bayes (BEB) and Naive empirical Bayes (NEB) methods were used, the amino acid sites under selection were quite variable and did not converge. Only the NEB sites were found to be significant in the two genes of interest, even though CODEML suggests that BEB results should be prioritized. However, among the 67 genes, there were no coloration genes found for vertical bar loss (Figure 5). Interestingly, the two coloration genes found under positive selection for vertical bar gain were related to iridophores and melanophores, respectively.

**Figure 5:**
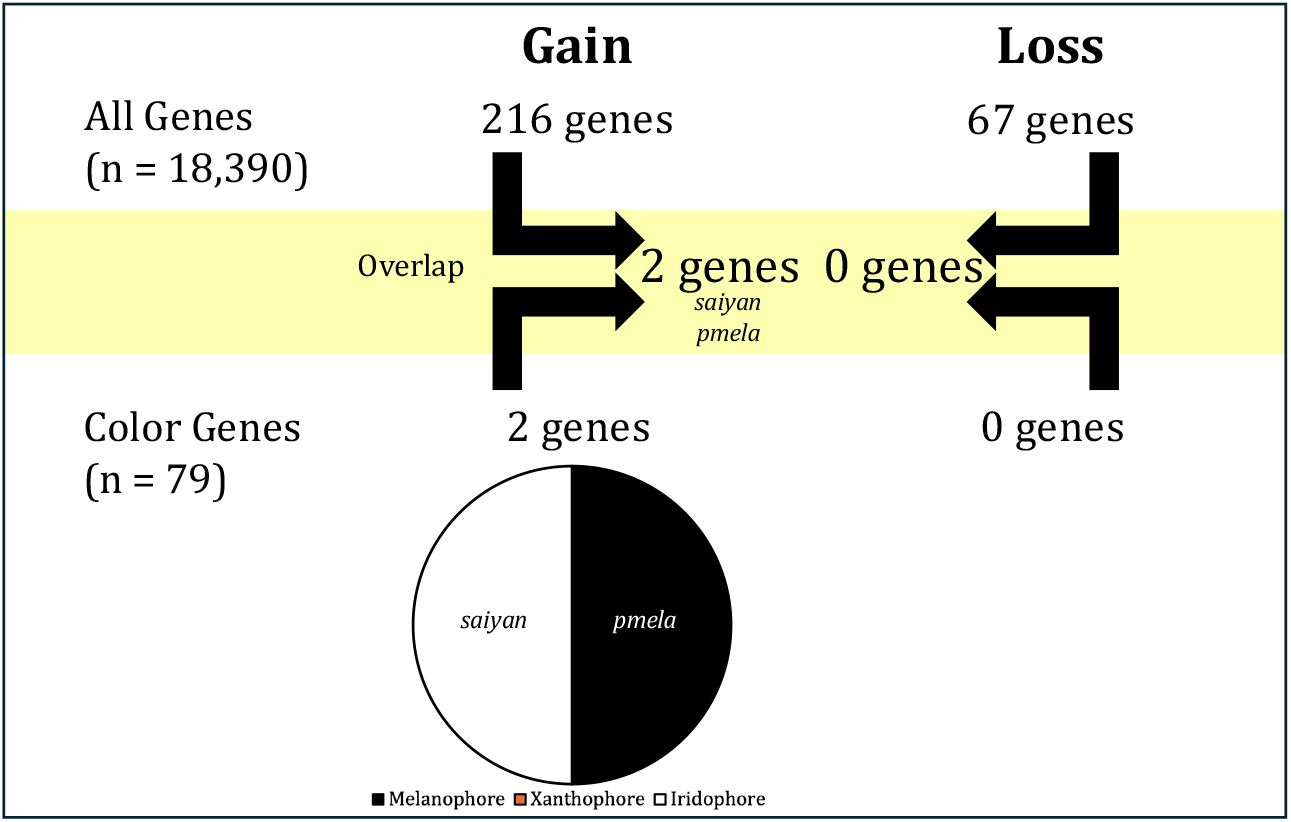
Summary of the number and name of significant color genes found in each dataset (All Genes and Color Genes) for gain and loss and their pigment cell type.

We found only three significant genes (*traf4a, adgrd1* and an unknown gene) that overlapped between the *BayesCode* gain analysis (n = 164) and CODEML gain analysis (n = 216), and none of them are linked to coloration. No genes were found between *BayesCode* loss analysis and CODEML loss analysis.

## Discussion

In this study, we investigated the genomic mechanisms underlying the evolutionary gain or loss of vertical bars in clownfish. Given previous ancestral state reconstruction in clownfish (Merilaita and Kelley, 2018; Salis et al., 2018), we expected that transitions in vertical bar patterns would be non-random and play a significant role in the evolutionary history of the group. Specifically, shifts in vertical bar presence or absence could be associated with ecological factors such as habitat complexity, social signaling, or changes in predation pressure. Based on this, we hypothesized that coloration genes involved in bar formation and maintenance would be under selection, either due to functional constraints maintaining the pattern or adaptive pressures favoring its modification. Our findings support this hypothesis, revealing that pigmentation-related genes exhibit signatures of positive selection and changes in selection pressure.

Our results of the ancestral state reconstruction revealed a high probability that the root of the clownfish clade had three vertical bars in line with previous studies (Salis et al., 2018). Vertical bar gain and loss appears to have occurred multiple times throughout clownfish evolution, with equal numbers of transitions in both directions. This is particularly interesting since prior studies (Merilaita and Kelley, 2018; Salis et al., 2018) have primarily focused on vertical bar loss rather than gain, whereas our results suggest that both processes are integral to clownfish coloration evolution. The repeated loss and gain of vertical bars across independent lineages coincides with shifts in ecological specialization, particularly in reproductive host associations. Recent work has shown that clownfish specializing in different sea anemone hosts exhibit convergent color patterns, with generalists typically retaining three white bars, whereas specialists tend to have reduced number of bars or altered pigmentation (Gaboriau et al., 2024). This pattern suggests that transitions in vertical bars are not random but instead linked to host-driven selection pressures, where specialists experience relaxed selection on disruptive coloration or adaptive shifts favoring different visual signals. Our findings reinforce this idea, demonstrating that vertical bar transitions have played a dynamic role in the evolutionary history of clownfish, potentially shaped by their ecological niches and host associations.

Since vertical bars transitions occurred multiple times in the clownfish evolution, we investigated if these changes were associated with shifts in selection. Using a Bayesian framework, we tested for signs of changes in selective pressure on pigmentation-related genes and genes that could be related to pigmentation. Our analysis identified several coloration genes under increased rate of evolution associated with vertical bar gain. The range of the Δ*ω* is quite small with all of the values well below 0.5 suggesting small effect size. These genes are linked to all three pigment cell types found in clownfish: melanophores, xanthophores, and iridophores. While Salis et al. (2021) provided a candidate list of coloration genes, we found specific examples that link how these genes directly affect color patterns in fish. It has been shown that mutations in *gch2* alters the xathophore pigmentation in zebrafish larvae, but in adults, normal pigmentation is restored (Lister, 2019). Additionally, *slc2a15b* plays a role in xanthophore and leucophore differentiation in teleost fish (Kimura et al., 2014), and CRISPR mutations of *oca2* lead to albinism in cavefish by disrupting the melanin pathway (Klaassen et al., 2018). These findings suggest that reduced purifying selection may have permitted greater genetic variation in bar gain-related genes, potentially enabling the recurrent evolution of vertical bars in different lineages.

Conversely, genes associated with vertical bar loss under increased rate of evolution were varying in their Δ*ω* values, with three genes (*gchfr, drd2a*, and *adam17b*) having a very large Δ*ω* at a single branch (well exceeding 1), suggesting strong purifying selection. These genes were only found to be related to melanophores and xanthophores, which control black and orange pigmentation, respectively. One gene of interest, *vps11*, has been linked to melanophore development (Clancey et al., 2013) and visual function in zebrafish (Banerjee et al., 2022). Given the importance of vision in species recognition within *Pomacentridae* (Stieb et al., 2017; Cortesi et al., 2020), the association between vision-related genes and bar loss may reflect adaptations in visual signaling.

The two genes found under positive selection were exclusively associated with vertical bar gain, with no significant findings for vertical bar loss. This distinction suggests that while increased rate of evolution plays a role in facilitating bar loss, directional selection may be maintaining or facilitating the re-emergence of bars in certain lineages. Although they are both significant, the amino acid sites under selection were quite variable and not convergent. None of the coloration genes under increased rate of evolution were found under positive selection, which is consistent with the idea that such changes in selective allows for accumulation of neutral mutations, whereas positive selection favors functional modifications. Given that the genes under positive selection were exclusively linked to vertical bar gain, we examined their potential functional roles in pigmentation. Interestingly, *saiyan* (*si:ch211-256m1*.*8*) was described by Salis and colleagues as significantly expressed in white skin cells of *A. ocellaris* and *A. frenatus* with no previous associated function (Salis et al., 2019b). Another gene, *pmel*, is essential for melanogenesis and has been shown to influence pigmentation in zebrafish and Nile tilapia, impacting melanophore and xathophore size and number (Wang et al., 2022). CRISPR gene editing confirmed that *pmela* also plays an important role in pigmentation, ocular structure and function (Lahola-Chomiak et al., 2018). Additionally, *pmel* has also been implicated in pattern transition from blotches to stripes in snakes (Tzika et al., 2024).

Our analyses revealed limited overlap between the genes under changes in the rate of evolution and positive selection results. Only three genes (*traf4a, adgrd1* and an unknown gene) were significant in both the *BayesCode* gain analysis (n = 164) and CODEML gain analysis (n = 216), with none being directly linked to coloration. Interestingly however, *TRAF4a* is involved in immune function and cold-water adaptation in fish (Kim et al., 2023) potentially having played a role in environmental adaptation. Additionally, six genes overlapped between the CODEML gain and loss analyses, but these genes were not functionally annotated. These minimal overlaps highlight key methodological differences: *BayesCode* identifies changes in selective pressure along a branch across the whole gene, indicating reduced evolutionary constraint, whereas CODEML detects positive selection, indicating functional divergence at specific sites of the gene. Given these differences, a lack of overlap is expected, as they do not capture the same signal of selection acting on sites and genes. Although we identified many additional significant genes associated with vertical bar gain and loss beyond known coloration genes, no significant GO terms related to color, pigmentation, or vision were detected. However, this does not mean these genes are unrelated to coloration; rather, their roles may simply not be annotated as such in current databases.

Overall, our findings contribute to a broader understanding of how selection shapes phenotypic diversity and adaptation in marine organisms. We identify several coloration genes under selection, which have been proven to be functionally related to color in other species, highlighting their potential role in clownfish coloration. The next step in validating these candidate genes is functional testing via CRISPR/Cas9, which has been successfully applied in clownfish but remains limited in scope (Mitchell et al., 2021). Future studies integrating gene expression analyses with functional assays will be crucial for determining the mechanistic basis of vertical bar evolution in clownfish.

## Supplementary material

Supplementary material is available in file “SupplementaryFiles.xlsx”

## Data and resource availability

All code is available on Github https://github.com/phylolab/clownfishBarEvolution.git

## Acknowledgments

The work was funded by a grant from the Swiss National Science Foundation to NS (grant 315230 219757) and from funding from the University of Lausanne. We also thank the DCSR infrastructure of the University of Lausanne for the computing resources and the Genomic Technologies Facility of the UNIL for genome sequencing.

## Author contributions

Original idea: LF, TL, and NS

Code: LF, TL, AM, TG

Data analyses: LF, TL, AM, TG, NS

Interpretation: LF, TL, AM, TG, DAH, and NS

First draft: LF

Editing and revisions: LF, TL, AM, TG, DAH, and NS.

Project management and funding: NS

